# Detecting dipicolinic acid production and biosynthesis pathways in Bacilli and Clostridia

**DOI:** 10.1101/803486

**Authors:** Benjamin Gordon, Paul Duellman, Anthony Salvucci, Marthah De Lorme

## Abstract

Bacterial endospores are highly resistant structures and dipicolinic acid is a key component of their resilience and stability. Due to the difficulty in controlling endospore contaminants, they are of interest in clean rooms, food processing, and production industries, while benefical endospore-formers are sought for potential utility. Dipicolinic acid production has traditionally been recognized in Bacilli, Clostridia, and Paenibacilli. Here, sixty-seven strains of aerobic and anaerobic endospore-forming bacteria belonging to the genera *Bacillus, Brevibacillus, Clostridium, Fontibacillus, Lysinibacillus, Paenibacillus, Rummeliibacillus*, and *Terribacillus* were grown axenically and sporulated biomasses were assayed for dipicolinic acid production using fluorimetric detection. Strains testing positive were sequenced and the genomes analyzed to identify dipicolinic acid biosynthesis genes. The well-characterized biosynthesis pathway was conserved in 59 strains of Bacilli and Paenibacilli as well as two strains of Clostridia; six strains of Clostridia lacked homologs to genes recognized as involved in dipicolinic acid biosynthesis. Our results confirm dipicolinic acid production across different classes and families of Firmicutes. We find that members of *Clostridium* (cluster I) lack recognized dipicolinic acid biosynthesis genes and propose an alternate genetic pathway in these strains. Finally, we explore why the extent and mechanism of dipicolinic acid production in endospore-forming bacteria should be fully understood. We believe that understanding the mechanism by which dipicolinic acid is produced can expand the methods to utilize endospore-forming bacteria, such as novel bacterial strains added to products, for genes to create inputs for the polymer industry and to be better equipped to control contaminating spores in industrial processes.

## INTRODUCTION

Bacterial spores are highly resistant structures in a dormant state, with little, if any, detectable metabolic activity (McKenney *et al.* 2013) (Figure 1). Spores are formed in response to adverse environmental conditions and spore formation, or sporulation, generally occurs when bacteria are challenged by nutritional stress (Driks 2002; Errington 2003; Piggot and Hilbert 2004). Spores can remain alive for extended periods of time and possess the ability to reactivate if nutrients become available and conditions are favorable (Moir 2003; Setlow 2003; Paredes-Sabja *et al.* 2011). Spores can survive many conditions that would otherwise destroy vegetative cells. The unique architecture of spores, such as the spore coat, cortex, and core, explains in large part their resistance to stresses and their ability to survive under extreme conditions (Henriques and Moran 2007).

**Figure 1.**
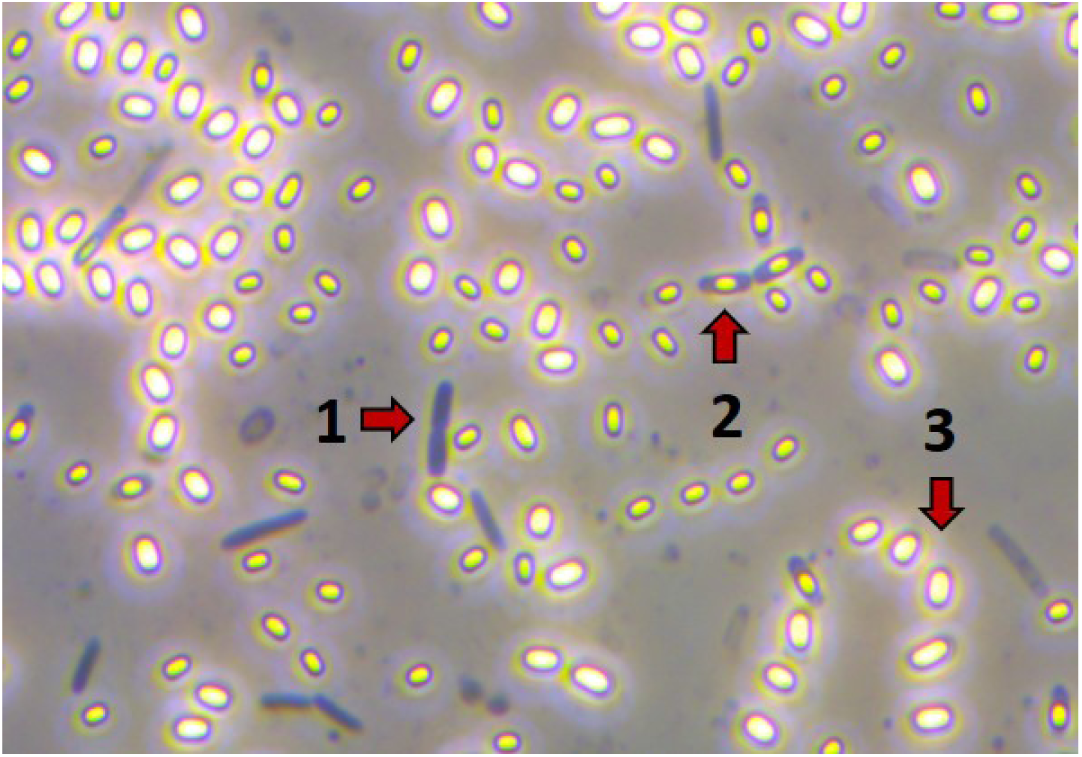
Phase-contrast microscopy (100X) of *Bacillus sp.* AS739 with examples of vegetative cells (1), endospores (2), and mature spores (3) visible.

Firmicutes are characterized by their ability to produce endospores (Figure 2-6), and compared to other spore-forms endospores are many times more resistant to oxidizing agents, heat, desiccation, and radiation (Setlow 1995). Pyridine-2,6-dicarboxylic acid, or dipicolinic acid (DPA) is an important component of bacterial endospores (Powell 1953). It has been shown that DPA is located in the spore core, and can represent 5–14% of endospore dry weight (Murrell 1969). DPA is maintained within intact endospores, and can be degraded under aerobic (Arima and Kobayashi 1962; Taylor and Amador 1988; Amador and Taylor 1990) and anaerobic (Seyfried and Schink 1990) conditions after it is released during spore germination. Dipicolinic acid forms a complex with calcium ions within the endospore core. This complex binds free water molecules causing dehydration of the spore. As a result, the heat resistance of macromolecules within the core increases (Gerhardt 1989). In addition, the calcium-dipicolinic acid complex also functions to protect DNA from heat denaturation by inserting itself between the nucleobases, thereby increasing the stability of DNA (Moeller *et al.* 2014).

**Figure 2.**
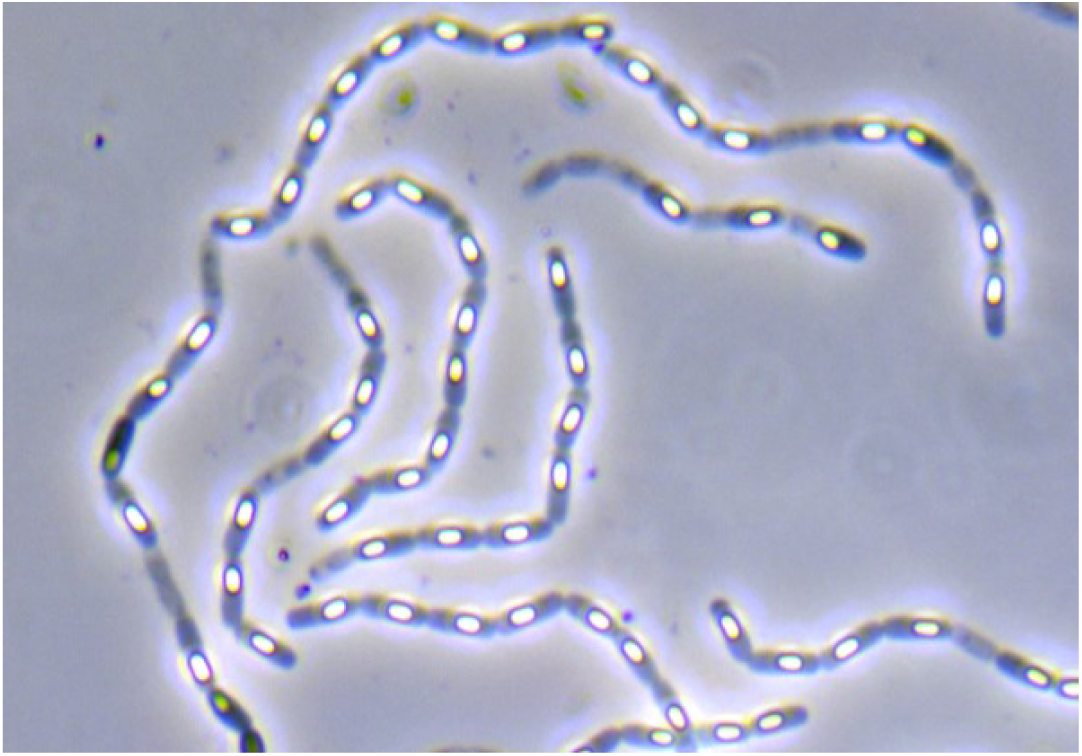
Phase-contrast microscopy (100X) of *Bacillus marisflavi* AS47 with endospores visible throughout.

**Figure 3.**
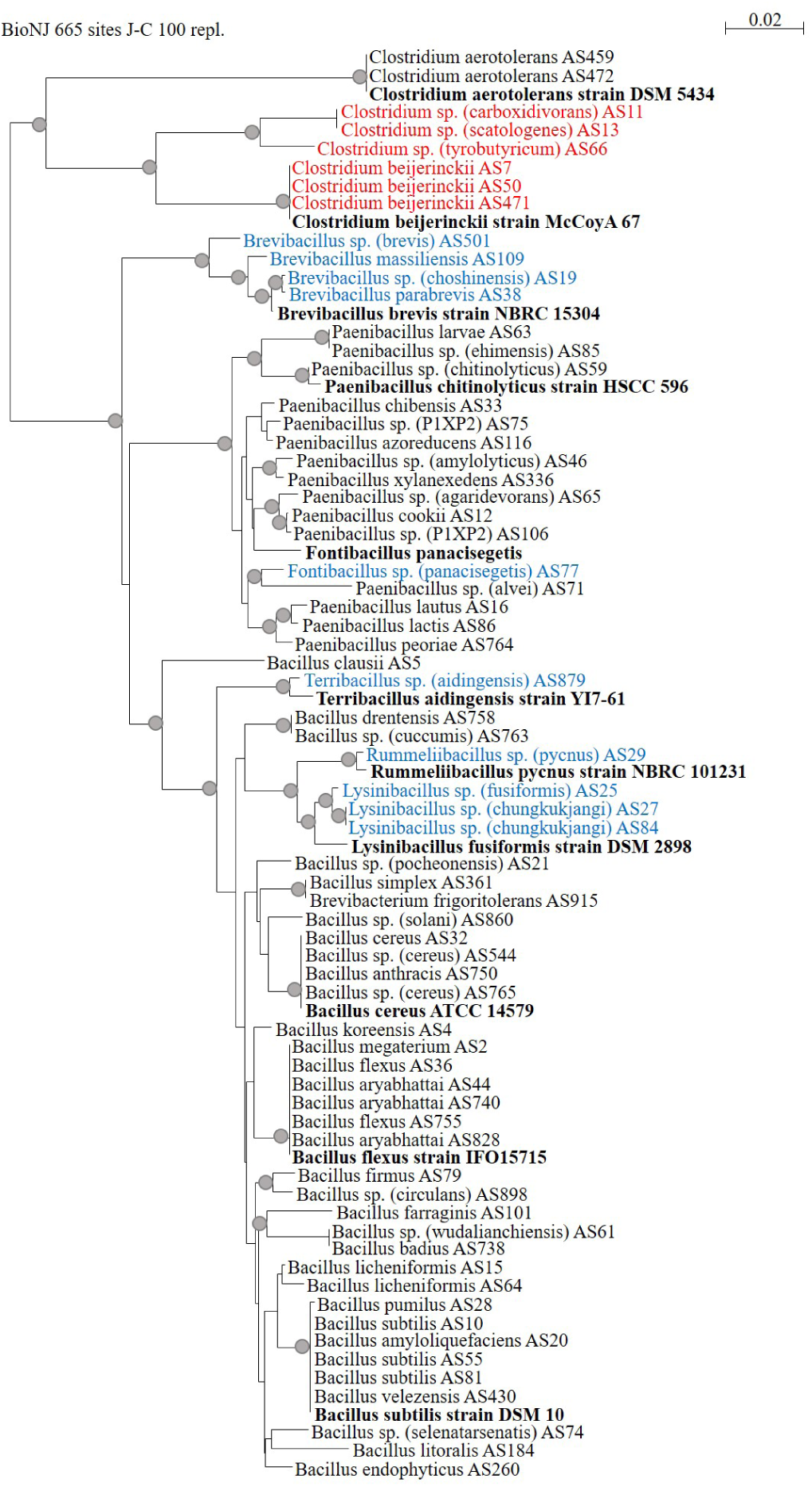
Neighbor-joining tree representing 16S rRNA-based phylogenetic distrtibution of strains assayed herein. Reference strains are highlighted in bold. Recently re-classified *Bacilli* and *Paenibacilli* experimentally demonstrated as producing DPA are highlighted in blue. Strains where known DPA synthase genes were not detectable are highlighted in red. Bootstrap support of >70% is notated by grey circles at branch points (Hillis and Bull 1993).

**Figure 4.**
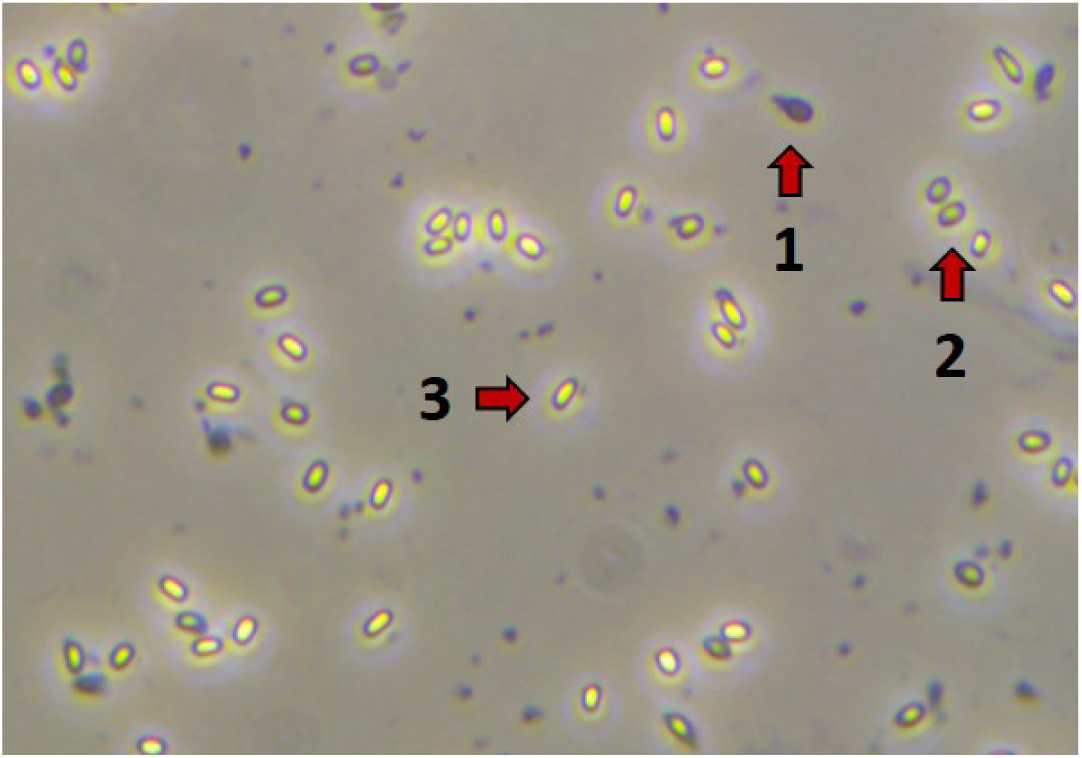
Phase-contrast microscopy (100X) of *Terribacillus sp. (aidingensis)* AS879 with examples of vegetative cells (1), endospores (2), and mature spores (3) visible.

**Figure 5.**
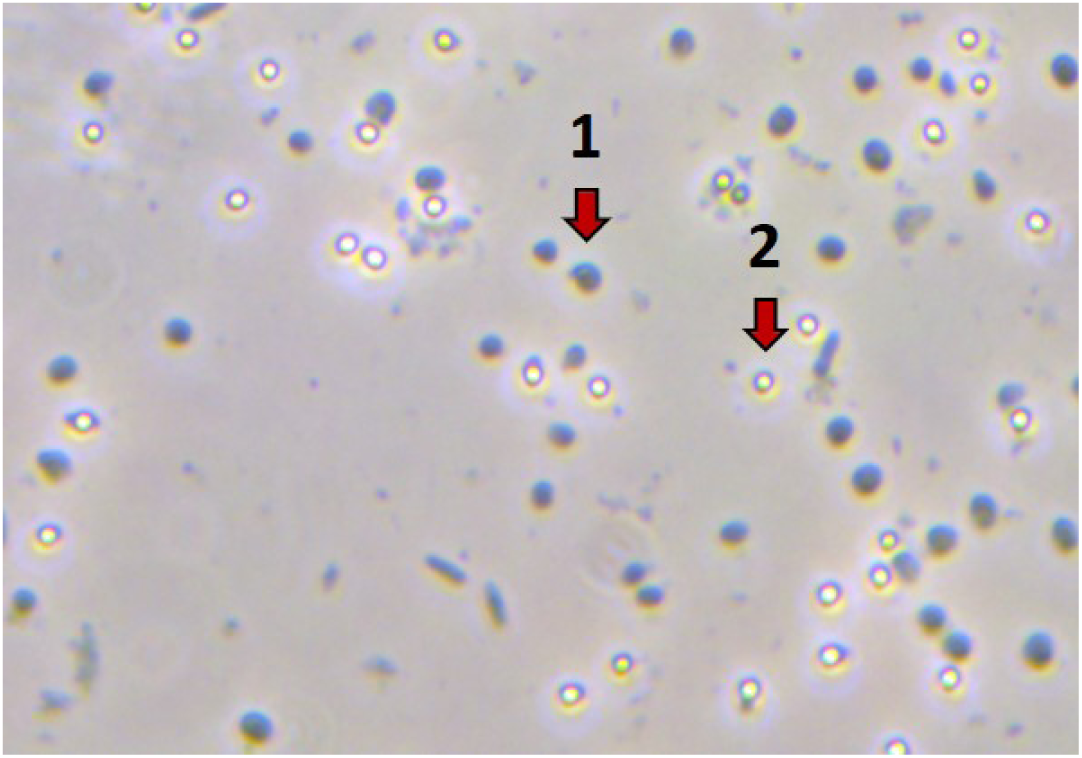
Phase-contrast microscopy (100X) of *Clostridium aerotolerans* AS472 with examples of vegetative cells (1) and mature spores (2) visible.

**Figure 6.**
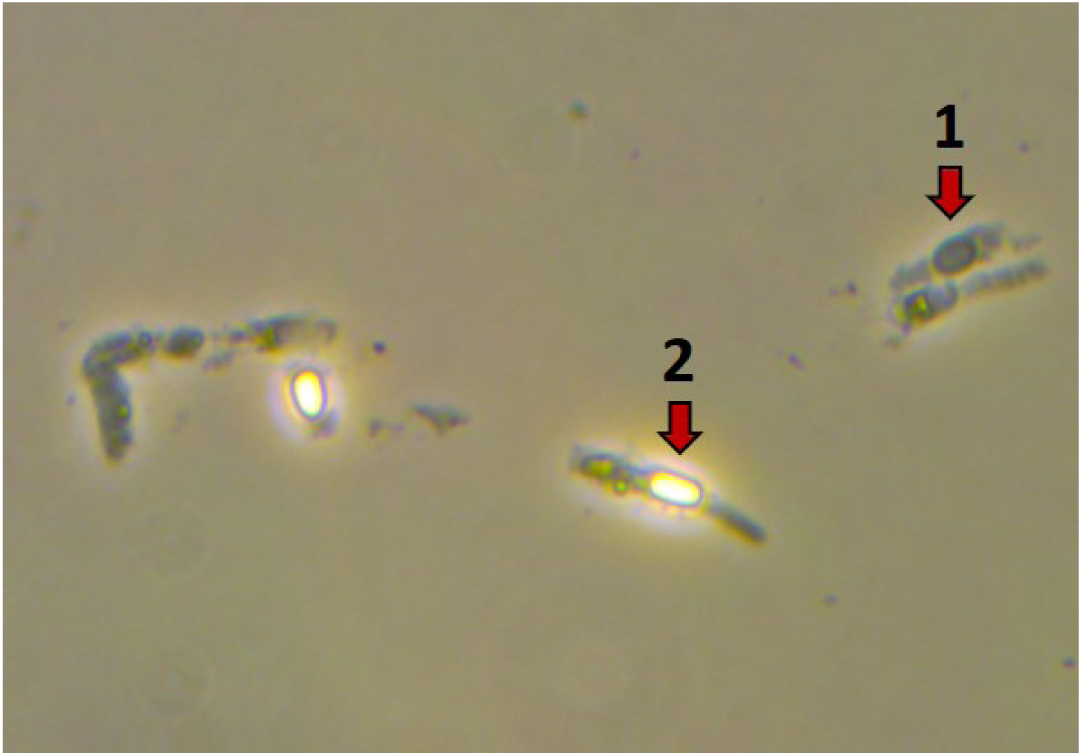
Phase-contrast microscopy (100X) of *Clostridium scatologenes* AS13 with examples of vegetative cells (1) and mature spores (2) visible.

In Bacilli, the DPA biosynthetic pathway has been well characterized (Wolska *et al.* 2007). In *B. subtilis*, the DPA synthetic pathway is encoded by four operons: *dapG, asd, dapA* and *dpaAB*. Asparatate kinase, encoded by *dapG*, is responsible for the first step of the biosynthesis cascade, producing L-4-aspartyl phosphate from L-aspartate. Aspartate-semialdehyde dehydrogenase, encoded by *asd*, is responsible for the second step, producing L-aspartate 4-semialdehyde. Dihydrodipicolinate synthase, encoded by *dapA* is responsible for the third step, producing L-2,3-dihydrodipicolinate (Takahashi *et al.* 2015). These steps are also used in lysine biosynthesis. DPA synthase which is responsible for the production of DPA, is encoded by *dpaAB*. Dipicolinate synthase subunit A (*dpaA*, otherwise known as *spoVFA*) encodes a putative dehydrogenase, and dipicolinate synthase subunit B (*dpaB*, otherwise known as *spoVFB*) appears to be a flavoprotein (Daniel and Errington 1993).

The major genera identified as endospore forming bacteria include *Bacillus, Paenibacillus*, and *Clostridium* (Fritze 2004; Logan and Halket 2011; Galperin 2013). Since 1990, the genus *Bacillus* has been split into several families and genera of endospore-forming bacteria based on 16S rRNA analysis (Galperin 2013). The unifying characteristic of these bacteria is that they are Gram-positive, form endospores, and aerobic. An increased concentration of DPA is a biochemical hallmark for endosporulating bacteria such as *B. subtilis* (Piggot and Hilbert 2004). We therefore hypothesized that strains capable of sporulating and previously classified as *Bacillus*, such as *Brevibacillus, Fontibacillus, Lysinibacillus, Rummeliibacillus*, and *Terribacillus* should produce DPA and contain the same set of genes responsible for DPA synthesis in Bacilli.

Work by others has shown that members of the *Clostridium sensu stricto* (cluster I) lack genes with significant homology to *dpaAB* (Stragier 2002; Onyenwoke *et al.* 2004) but nevertheless produce DPA during sporulation (Paredes-Sabja *et al.* 2008). This cluster is particularly important due to the presence of industrially useful strains such as *Clostridium acetobutylicum* and *C. beijerinckii*, as well as human pathogens such as *C. perfringens, C. botulinum* and *C. tetani*. Osburn et al. (2010) implicated an electron transfer flavoprotein *α*-chain (*etfA*) that is directly involved in DPA synthesis in *C. perfringens* using a modified version of the Bach and Gilvarg assay system (Bach and Gilvarg 1966; Orsburn *et al.* 2010). We hypothesized that other members of the *Clostridium* (cluster I) such as *C. beijerinckii, C. carboxidivorans, C. scatologenes*, and *C. tyrobutyricum*, use the same or similar biosynthesis pathways.

DPA can be detected by a range of analytical techniques (Beverly *et al.* 1996; Yilmaz *et al.* 2010; Cowcher *et al.* 2013; Wang *et al.* 2011). The terbium dipicolinate fluorescence method (Rosen *et al.* 1997; Hindle and Hall 1999) is both tractable and sensitive, allowing the detection of nanomolar concentrations. This technique is therefore appropriate for testing if the genera hypothesized to produce DPA (*Brevibacillus, Fontibacillus, Lysinibacillus, Rummeliibacillus*, and *Terribacillus*) do in fact have the capacity for DPA-production. The terbium dipicolinate fluorescence method can also be used to establish that Clostridia such as *C. beijerinckii, C. carboxidivorans, C. scatologenes*, and *C. tyrobutyricum* produce DPA prior to sequencing and genomic evaluation to look for *dpaAB* and *etfA* DPA biosynthesis gene homologues.

In the present study, bacteria isolated from soil and a microbial product were taxonomically identified using 16S rRNA sequence analyses. Presumptive endospore-forming bacteria were then screened for DPA production *in vitro*. DPA detection was followed by whole genome-sequencing to confirm taxonomic identifications at the species-level and to examine whether known DPA biosynthesis pathways were present.

## MATERIALS AND METHODS

### Strain Isolation and Initial Identification

All strains used were isolated from either bulk soil or from a sample of HYT® A (Table S1) (Agrinos, https://agrinos.com). Strains were then repeatedly streaked onto semi-solid media until isogenic. Initial taxonomic classifications were made using 16S rRNA gene sequences. Full length 16S genes were amplified from each strain using genomic DNA and/or colonies as the PCR template (27F, 5’-AGRGTTTGATCMTGGCTCAG-3’; 1492R, 5’-GGTTACCTTGTTACGACTT-3’; following (Singer *et al.* 2016) with minor modifications). PCR products were sequenced directly using the forward and reverse PCR primers. The high-quality sequence traces were then analyzed and a BLAST analysis was performed using the NCBI 16S ribosomal RNA sequence database (Madden 2013).

### Terbium Dipicolinate Fluorescence Assay

Prior to testing, strains were grown as follows. Aerobic strains were cultured on Trypticase Soy Agar (i.e. TSA; Tryptone 17 g/L, Soytone 3 g/L, Dextrose 2.5 g/L, Sodium chloride 5 g/L, Dipotassium phosphate 2.5 g/L, Agar 15 g/L) in a static incubator at 30° for up to three days. Anaerobic strains were cultured on Reinforced Clostridial Medium (i.e. RCM; BD Difco™ Reinforced Clostridial Medium, Cat No. 218081) in a static incubator at 35° for up to five days.

All cultures were visualized using phase-contrast microscopy at 100X magnification to verify the presence of refractile spores (Bulla *et al.* 1969). A modified version of the protocol used by Hindle and Hall (1999) was used to detect the presence of DPA. Using a sterile disposable loop, a 1 µl loopful was taken off a plate and resuspended in 10 ml of sodium acetate buffer (0.2 M, pH 5) in screw cap glass test tubes. Samples were autoclaved for 15 minutes at 121° to heat the spores and release any DPA. Samples were then cooled for 10 min in a room temperature (22°) water bath. Following cooling, for each sample 100 µl was mixed with 100 µl of terbium chloride solution (TbCl3, 30 µM) in wells of a 96-well flat-bottomed clear microtiter plate. Fluorescence was measured with a BioTek® Cytation 5 Multi-Mode Reader (BioTek Instruments, Inc., https://www.biotek.com/) using time resolved fluorescence (delay 50 µs, excitation wavelength 272 nm, emission wavelength 540 nm) (Brandes Ammann *et al.* 2011). Strains testing higher than negative controls, with a fluorescence intensity of more than 10,000 relative fluorescence units, and endospores visible with phase-contrast microscopy were considered DPA-producers.

### Whole-genome sequencing and Bioinformatic Analysis

Whole-genome sequencing was performed by CoreBiome (Core-Biome, Inc., https://www.corebiome.com). Briefly, isolates were grown in appropriate liquid culture. A minimum of 1 × 10^9^ cells were spun down and the supernatant removed prior to freezing at −20°. Samples were then sent to CoreBiome for DNA extraction, followed by whole-genome sequencing, assembly, and annotation using their StrainView™ whole-genome shotgun sequencing service. Genome assembly quality was assessed using QUAST (Gurevich *et al.* 2013). Whole-genome sequence annotation was performed using Prokka (https://github.com/tseemann/prokka) (Seemann 2014). 16S sequences were identified using Barrnap (https://github.com/tseemann/barrnap). To assign function to CDS features, a core set of well characterized proteins are first searched using BLAST+, then a series of slower but more sensitive HMM databases are searched using HMMER3 (http://hmmer.org) (Finn *et al.* 2011).

### Phylogenetics

For each sequence, a BLAST analysis was performed using the NCBI 16S ribosomal RNA sequence database (Madden 2013). To make species-level identifications, representative genomes were retrieved from NCBI and the whole-genome similarity metric Average Nucleotide Identity (ANI) was calculated for top 16S BLAST hits. Whole-genome comparisons with ANI values of greater than 95% were considered to be the same species (Konstantinidis and Tiedje 2005; Goris *et al.* 2007). For whole-genome comparisons with ANI values less than 95%, strains were labeled as *Genus sp. ()*, with the closest known relative indicated. Using Seaview Version 4.6.1 (Gouy *et al.* 2009; Galtier *et al.* 1996), 16S sequences were aligned using MUSCLE (Edgar 2004) and BioNJ trees (Gascuel 1997) were computed using Jukes & Cantor (Jukes and Cantor 1969) pairwise phylogenetic distances with all gap containing sites excluded from the analysis.

### Data Availability

Table S1 contains a list of all strains used in this study, RFUs measured using the terbium-DPA fluorescence assay, and DPA biosynthesis gene copy-number for each strain. File S1 contains an alignment of 16S rRNA sequences. File S2 contains an alignment of *dpaA* protein sequences. File S3 contains an alignment of *dpaB* protein sequences. File S4 contains an alignment of *Isf* protein sequences.

## RESULTS

Of the 67 strains included in this study (Table S1), all were Firmicutes representing the classes of Bacilli, Clostridia and Paenibacilli. As expected, all 49 strains of *Bacillus* and *Paenibacillus* (Figure 3, in black) were found to form endospores via phase-contrast microscopy (see Figures 1 and 2). DPA biosynthesis was detected *in vitro* (Table S1), and genome sequences revealed that all of these strains possessed known DPA synthase genes (Files S2 & S3).

Ten strains from five genera that were re-classified from *Bacillus* to *Brevibacillus, Fontibacillus, Lysinibacillus, Rummeliibacillus*, and *Terribacillus* which were hypothesized to produce DPA (Figure 3, highlighted in blue) were found to form endospores via phase-contrast microscopy (see Figure 4). DPA biosynthesis was detected *in vitro* (Table S1), and genome sequences revealed that all of these strains possessed known DPA synthase genes, *dpaAB* (Files S2 & S3).

Eight strains of Clostridia which were expected to produce DPA were found to form endospores via phase-contrast microscopy (see Figures 5 and 6). DPA biosynthesis was detected *in vitro* (Table S1), and genome sequences revealed that two strains of *Clostridium aerotolerans* possessed homologs to known DPA sythase genes, *dpaAB* (Files S2 & S3). Genome sequences from the remaining six Clostridia (Figure 3, highlighted in red) did not have detectable known *dpaAB* or *etfA* genes (Table 1). *etfA* contains a flavin mononucleotide (FMN), therefore a search was performed for other flavoproteins that contained a FMN and were common among the six Clostridia that lacked homologs to known DPA synthesis genes.

**Table 1.** DPA biosynthesis gene copy count and levels of DPA detected in strains of *Clostridium*.

| Taxonomy<sup>a</sup> | *dpaA*<sup>b</sup> | *dpaB*<sup>c</sup> | *etfA*<sup>d</sup> | *Isf*<sup>e</sup> | RFU<sup>f</sup> |
| --- | --- | --- | --- | --- | --- |
| *Clostridium beijerinckii* AS471 | 0 | 0 | 0 | 10 | 103,839 |
| *Clostridium beijerinckii* AS50 | 0 | 0 | 0 | 11 | 386,574 |
| *Clostridium beijerinckii* AS7 | 0 | 0 | 0 | 6 | 65,340 |
| *Clostridium sp. (carboxidivorans)* AS11 | 0 | 0 | 0 | 1 | 48,558 |
| *Clostridium sp. (scatologenes)* AS13 | 0 | 0 | 0 | 1 | 45,462 |
| *Clostridium sp. (tyrobutyricum)* AS66 | 0 | 0 | 0 | 1 | 47,567 |

<sup>a</sup> For strains denoted as sp., the species in parenthesis is the closest recognized species <sup>b</sup> Dipicolinic acid synthase subunit A <sup>c</sup> Dipicolinic acid subunit B <sup>d</sup> Electron transfer flavoprotein $\alpha$ -chain <sup>e</sup> Iron-sulfur flavoprotein <sup>f</sup> Relative fluorescence units

While the *Clostridium* (cluster I) strains lack the recognized DPA biosynthesis genes, they all possess at least one copy of an iron-sulfur flavoprotein (*Isf*) (File S4), and strains with more *Isf* copies produced more DPA (Table 1). Both *etfA* and *Isf* are thought to be involved in electron transport (Tsai and Saier Jr 1995; Becker *et al.* 1998). It is possible that these two genes have analogous activity, catalyzing the formation of DPA from L-2,3-dihydrodipicolinate (Figure 7). Future work including *Isf* gene knockout mutants that lack DPA production and/or observing upregulation of *Isf* expression during sporulation would be consistent with *Isf* having a role in DPA formation.

**Figure 7.**
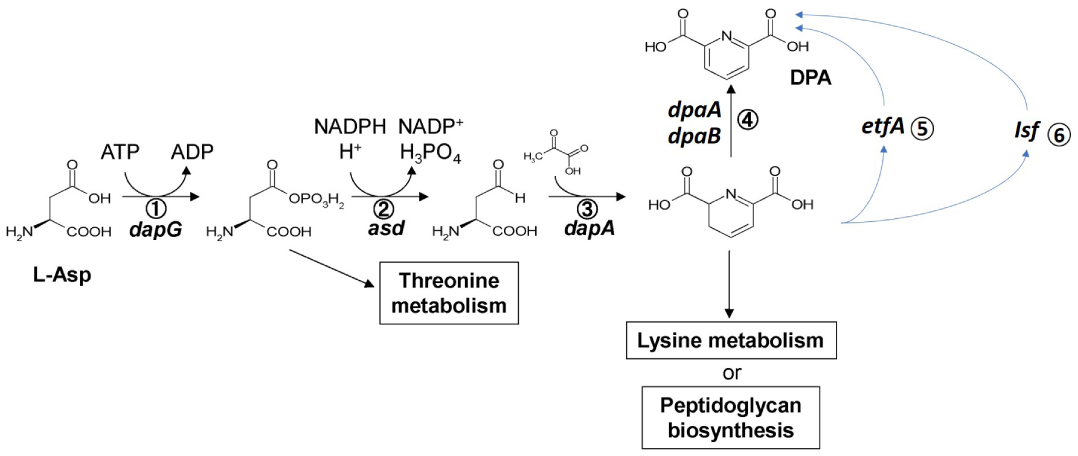
Dipicolinic acid biosynthesis pathways. ^1^*dapG*: aspartokinase; ^2^*asd*: aspartate-semialdehyde dehydrogenase; ^3^*dapA*: dihydrodipicolinate synthase; ^4^*dpaAB*: dipicolinate synthase. ^5^*etfA*: elctron transfer flavoprotein subunit A; ^6^*Isf*: iron-sulfur flavoprotein. Modified from (Takahashi *et al.* 2015).

## DISCUSSION

Here we have found that genera was a good predictor of DPA production, and added experimental evidence to support the production of DPA by additional Bacillaceae. This result, which is consistent with their previous classification as *Bacillus*, is yet an important observation given the functional diversity that these genera possess. In addition, we found the *Clostridium* (cluster I) lack recognized biosynthesis pathways for DPA production. This result, while similar to previous work (Orsburn *et al.* 2010), fails to confirm their findings and we propose an alternate genetic pathway in these strains.

Members of the Bacillaceae family found to produce DPA have recently been shown to possess many functions and utilities, and the survivability that DPA production adds, makes these strains ripe for use in novel products. For example, strains of *Brevibacillus* have been found to have larvicidal activity (Zubasheva *et al.* 2010), biological control of plant pathogens (Chandel *et al.* 2010; Liu *et al.* 2010), as well as improving shelf-life of fruits (Che *et al.* 2011). *Fontibacillus* strains have been shown to produce a novel and highly reusable pullanase (Alagöz *et al.* 2016) which is used as a processing aid for the production of ethanol or sweeteners from grain. *Lysinibacillus* species have been shown to degrade harmful insecticides (Singh *et al.* 2012), selectively desulfurize dibenzothiophene (Bahuguna *et al.* 2011), and produce mosquito larvicidal toxins (Lozano *et al.* 2011). In addition, strains can produce 10-hydroxystearic acid from olive oil (Kim *et al.* 2012). *Rummeliibacillus* species have been shown to transform palm oil mill effluent to polyhydroxyalkanoate and biodiesel (Junpadit *et al.* 2017), enhance growth and health in tilapia (Yih *et al.* 2019), and has the potential for biomineralization (Mudgil *et al.* 2018). Finally, *Terribacillus* species have been found to produce a novel chitinase as well as antifungal enzymes that repress plant diseases (Essghaier *et al.* 2014). Given the wealth of functionalities that this family possesses, coupled with a long shelf-life and potential to survive in a variety of manufacturing processes due to DPA production, makes Bacillaceae well suited for industrial processes and an untapped well of biotechnology potential.

We have confirmed here that some Clostridia possess a DPA synthase homolog. We have also confirmed that *Clostridium* (cluster I) do not possess a DPA synthase homolog. Unlike previous findings, we were unable to identify *etfA* in our assayed *Clostridium* (cluster I) strains. Despite the lack of *dpaAB* and *etfA* genes, these strains were still able to produce detectable quantities of DPA.

We propose the *Isf* gene product as an alternate pathway for DPA-production in *Clostridium* (cluster I). This alternate biosynthesis pathway for DPA synthesis represents a new potential route for bioengineering the production of DPA in other strains of bacteria via the addition of a single gene to the lysine pathway, particularly in anaerobes. Dipicolinic acid production has been engineered in bacterial strains using the traditional *dpaAB* genes (Zelder *et al.* 2009; McClintock *et al.* 2018). DPA has numerous industrial uses, for example, the monomer can be used as a replacement for petroleum derived isophthalic acid in the synthesis of polyesters or polyamide copolymers (McClintock *et al.* 2018) resulting in a more biodegradable compound. The scalable synthesis of DPA using a variety of bacteria could increase its use as a replacement for other aromatic dicarboxylic acids.

Probiotics have been developed for human use as well as for use in animal feeds, and a large number of these formulations utilize spore-forming *Bacillus* (Hong *et al.* 2005). Effective probiotics need to thrive in anaerobic conditions and retain viability in aerobic conditions on the shelf, as well as cross the gastric barrier and enter the upper GI tract in a viable state following consumption. Probiotics utilizing DPA-producing endospores are proven to have better survivability during passage through the acidic stomach environment, show better survival during manufacturing, and have a longer shelf-life (Bader *et al.* 2012). The potential of Clostridia as probiotics has only recently been discussed (Cartman 2011). DPA-containing Clostridial endospores are ideal vehicles for overcoming the challenges probiotic bacteria encounter, as they can survive aerobic storage and remain viable while crossing the host gastric barrier, followed by germination in the anaerobic environment of the upper GI tract where they are well adapted to survive. *Clostridium* have been documented as members of healthy GI tracts in humans (Arumugam *et al.* 2011; Qin *et al.* 2010; Tap *et al.* 2009), have been shown to increase resistance against Irritable Bowel Disease (Atarashi *et al.* 2011), and can protect against *C. difficile* infection (Merrigan *et al.* 2009; Sambol *et al.* 2002). Given the survivability of DPA-producing Clostridia, and the recently highlighted benefits of *Clostridium* as members of the gut microbiota, they may prove to be an ideal probiotic.

In addition, an alternate biosynthesis pathway for DPA represents a new target for designing drugs to prevent contamination, as inhibiting the production of DPA would make endospores more susceptible to traditional manufacturing procedures. Many of the endospore-forming bacteria can survive food processing, decreasing the shelf-life of processed foods (e.g. *Clostridium tyrobutyricum*), and potentially making them unsafe to consume (e.g. *Clostridium botulinum*) (Daelman *et al.* 2013; Soni *et al.* 2016; André *et al.* 2017). Endospore-forming *Clostridium* also contain members that are known for their pathogenic potential, such as *Clostridium* (e.g. (*C. perfringens, C. novyi, C. septicum, C. tetani*, and *C. difficile*) (Ehling-Schulz *et al.* 2019; Wells and Wilkins 1996).

Here we have added experimental evidence to support the production of DPA by additional Bacillaceae and we have proposed an alternate pathway in *Clostridium* (cluster I) strains that lack recognized dipicolinic acid biosynthesis genes. By fully understanding the breadth and mechanisms by which DPA is produced, we can utilize endospore-forming bacteria in new ways: as novel bacterial strains added to products, for genes to create inputs for the polymer industry, and to be better equipped to control contaminating spores in industrial processes.

## ACKNOWLEDGEMENTS

The authors would like to thank Frederic Kendirgi for his guidance during the research process. We are grateful to Chynna Gabotero, Mei Konishi, and Nicole Hensley for critical reading of the manuscript ahead of publication, and to Mylavarapu Venkatramesh for supervision during the study. This work was supported by Agrinos, and we would like to thank Agrinos management for their ongoing financial support.

